# The Case for Retaining Natural Language Descriptions of Phenotypes in Plant Databases and a Web Application as Proof of Concept

**DOI:** 10.1101/2021.02.04.429796

**Authors:** Ian R. Braun, Colleen F. Yanarella, Jyothi Prasanth Durairaj Rajeswari, Diane C. Bassham, Carolyn J. Lawrence-Dill

## Abstract

Similarities in phenotypic descriptions can be indicative of shared genetics, metabolism, and stress responses, to name a few. Finding and measuring similarity across descriptions of phenotype is not straightforward, with previous successes in computation requiring a great deal of expert data curation. Natural language processing of free text descriptions of phenotype is often less resource intensive than applying expert curation. It is therefore critical to understand the performance of natural language processing techniques for organizing and analyzing biological datasets and for enabling biological discovery. For predicting similar phenotypes, a wide variety of approaches from the natural language processing domain perform as well as curation-based methods. These computational approaches also show promise both for helping curators organize and work with large datasets and for enabling researchers to explore relationships among available phenotype descriptions. Here we generate networks of phenotype similarity and share a web application for querying a dataset of associated plant genes using these text mining approaches. Example situations and species for which application of these techniques is most useful are discussed.

**Database URLs:** The database and analytical tool called QuOATS are available at https://quoats.dill-picl.org/. Code for the web application is available at https://git.io/Jtv9J. Datasets are available for direct access via https://zenodo.org/record/7947342#.ZGwAKOzMK3I. The code for the analyses performed for the publication is available at https://github.com/Dill-PICL/Plant-data and https://github.com/Dill-PICL/NLP-Plant-Phenotypes.

## 1 INTRODUCTION

Phenotypes, defined as measurable characteristics or properties of an organism that result from interactions between genetics and the environment, comprise an enormous portion of the biological data considered important across a wealth of domains in the life sciences and beyond. Phenotypes are everything we see or measure in biology. On a more practical note, phenotypes encompass critical information related to human health and medicine, and important agronomic traits such as plant height and biomass of crop species. The scope of phenotypic information also ranges widely, from cellular phenotypes such as membrane composition or chemical concentrations, to community-level phenotypes like total leaf surface area in a field of crops. The extreme diversity in how phenotypes can be observed and represented makes handling this information on a computational level fundamentally different than genomic data, which lends itself to computational means of representation and analysis based on the existing natural codes of bases and amino acids (reviewed in Braun et al. 2018). This is especially true for phenotypes that are comparative, such as two different leaf morphologies, rather than phenotypes that are easily translated into a quantitative value, such as height (reviewed in Yanarella et al. 2020).

Despite these challenges, bio-ontologies have greatly helped to enable computation on phenotypic information by providing standardized, hierarchical sets of descriptors (terms) that can be used to annotate phenotypic information. Doing so enables comparison between data points, including comparisons across multiple species, studies, and sources in a meaningful way, which has contributed to the use of these data structures in recent years. Using terms from the Gene Ontology (GO; Ashburner et al. 2000) to describe cellular components, functions, and processes allows researchers to quickly find genes related to a biological concept of interest, and to understand which biological processes are potentially carried out or influenced by a group of genes of interest (Huang et al. 2009). Using this same ontology as the format for predictions about gene functions allows datasets of predicted gene functions to be seamlessly incorporated with and compared to known information (Zhou et al. 2019). Biomedical vocabulary graphs such as the Human Phenotype Ontology (HPO; Robinson et al. 2008) and Disease Ontology (DO; Schriml et al. 2012) allow for organization and interoperability of the vast and growing body of knowledge surrounding human medicine. Efforts such as Phenoscape (Edmunds et al. 2015), the Monarch Initiative (Mungall et al. 2017), and Planteome (Cooper et al. 2018), use ontologies to provide common data representations and allow for comparisons across diverse species or across evolutionary history.

At the same time, both the performance and availability of natural language processing (NLP) and machine learning (ML) methods for working with natural language and text data have continually improved. Large language models now include artificial intelligence (AI) tools such as ChatGPT, developed by OpenAI (OpenAI 2023). The release of ChatGPT is popular, in part, because of its public availability and conversational nature. ChatGPT uses a generative pre-trained transformer (GPT) model (Radford et al. 2018), a deep-learning language model, and has elicited an excited response within the language processing community (Dwivedi et al. 2023). Improvements in language processing are due to recent and continued innovations in how neural networks are designed to handle this type of information (Mikolov et al. 2013; Le and Mikolov 2014; Vaswani et al. 2017), and how they can be trained on massive volumes of unlabeled data (such as Wikipedia or PubMed) to provide systems for accurately modeling text in computable formats, and allowing for transfer to other domains and fine-tuning for more specific problems (Devlin et al. 2018, Wolf et al. 2020). One result of this progress is that such techniques now represent a complementary approach to computationally handle the diversity of phenotypic information, at least for cases where phenotypes are represented as text descriptions. Given that phenotypes have been described in academic articles for more than a century, sources for phenotypic descriptions abound. Although the vast majority of phenotypes described in the literature have not been extracted and represented in computationally accessible community databases, some databases do exist that contain phenotype descriptions in free text fields.

Previously, we demonstrated that for some organizational tasks (like grouping functionally similar genes together), computational approaches that process text descriptions of phenotypes can work as well as, or better than, curated ontology term annotations for the creation of meaningful similarity measurements (Braun and Lawrence-Dill 2019). Here, we demonstrate that this finding holds true for a larger dataset of the available phenotype text descriptions from across six different plant species. This means that, where available, text descriptions of phenotypes have the potential to provide useful biological insight when combined with a variety of methods from the field of NLP. We therefore make a case for expanded inclusion of free text descriptions as a valuable component of biological databases going forward, whether as a supplemental data type to more standardized ontology term annotations, or as a potential short-term alternative for species currently lacking the curatorial resources to produce large scale datasets of high-confidence, curated annotations.

In demonstrating the utility of analyzing text descriptions of phenotypes with NLP approaches, we focus on what can be learned from evaluating similarity between descriptions as a measure of gene pair similarity. This is closely comparable to the ongoing problem in NLP of measuring sentence similarity, which has applications for text querying, text classification, and other tasks (De Boom et al. 2016). An enormous variety of solutions have been put forward for this problem, including both general solutions as well as more narrowly focused solutions for working in particular domains, such as biomedical literature (Soğancıoğlu et al. 2017, Chen et al. 2019). The number of solutions to this task is related to the fact that virtually all approaches for dealing with text computationally involve representing words or sentences as numerical vectors, on top of which similarity or distance metrics can then be applied to quantify relatedness between the two texts. In other words, all approaches for vectorizing text, which is typically the first step in handling any problem with NLP, can subsequently be used to find similarity between two texts by applying similarity metrics to their vector representations. This enables the generation of networks for organizing data across large datasets. In this work, we assess the performance of a variety of both simple and state-of-the-art methods for translating plant phenotype descriptions into numerical vectors and build networks that can be used to make inferences from pairwise similarities.

We also discuss and demonstrate how these same techniques can be applied for organizing and analyzing large phenotype description datasets, accounting for phenotypic characteristics that have not yet been explicitly defined by the input data. Finally, we provide a web application that enables others to explore and make use of phenotypic similarities identified. The application, called QuOATS (Querying with Ontology Annotations and Text Similarity), can be used to search for plant genes with similar phenotypic descriptions using gene identifiers, ontology terms, keywords, or similarity to searched phenotype descriptions as input.

## 2 MATERIALS AND METHODS

### 2.1 Datasets

Species included for our analyses included *Arabidopsis thaliana* (L.) Heynh. (Arabidopsis), *Zea mays* L. subsp. *mays* (maize), *Medicago truncatula* Gaertn. (barrel medic or Medicago), *Oryza sativa* L. (rice), *Glycine max* (L.) Merr. (soybean), and *Solanum lycopersicum* L. (tomato). We collected a dataset of available phenotype descriptions that have been mapped to specific plant genes, primarily through mutation studies, from the model species databases for Arabidopsis - TAIR (Berardini et al. 2015), maize - MaizeGDB (Portwood et al. 2019), and solanaceous plants - SGN (Fernandez-Pozo et al. 2015), and combined these data with as a dataset of phenotype descriptions created by Oellrich et al. 2015 that includes all six species. After merging data from multiple sources and preprocessing the texts, the combined dataset consisted of 7,907 genes from the 6 plant species, with the quantity of genes and the text describing their associated phenotypes varying across species (Table 1). The distributions of sentences and words quantities present per gene also vary broadly across species (Figure 1). Portions of the vocabulary used to describe phenotypes in each of the species are unique to that particular species, but in all cases more than 80% of the vocabulary was shared with at least one other species (Figure 2).

**Table 1.** Scope and scale of the complete dataset.

| Species | Number of Genes | Phenotype Descriptions |  |  |  | Annotations |  |  | Other Databases |  |  |  |
| --- | --- | --- | --- | --- | --- | --- | --- | --- | --- | --- | --- | --- |
|  |  | Total Sentences | Total Words | Unique Words | Unique to Species | Mapped to EQ Stmt(s) | Mapped to GO Term(s) | Mapped to PO Term(s) | Mapped in PlantCyc | Mapped in KEGG | Mapped in STRING | Mapped in PANTHER |
| Arabidopsis | 6,274 | 30,123 | 261,422 | 6,792 | 5,084 | 2,393 | 5,984 | 4,395 | 843 | 1,435 | 4,317 | 377 |
| Maize | 1,405 | 7,512 | 47,139 | 1,722 | 498 | 114 | 190 | 761 | 133 | 157 | 229 | 443 |
| Rice | 92 | 478 | 3,689 | 760 | 97 | 92 | 46 | 92 | 3 | 0 | 45 | 86 |
| Tomato | 69 | 359 | 1,678 | 552 | 99 | 72 | 25 | 64 | 2 | 18 | 11 | 10 |
| Medicago | 37 | 263 | 2,447 | 671 | 123 | 40 | 30 | 32 | 2 | 0 | 13 | 0 |
| Soybean | 30 | 62 | 222 | 78 | 12 | 30 | 28 | 27 | 0 | 1 | 5 | 0 |
| Total | 7,907 | 38,797 | 316,597 | 7,663 | 5,913 | 2,741 | 6,303 | 5,371 | 983 | 1,611 | 4,620 | 916 |

**Figure 1.**
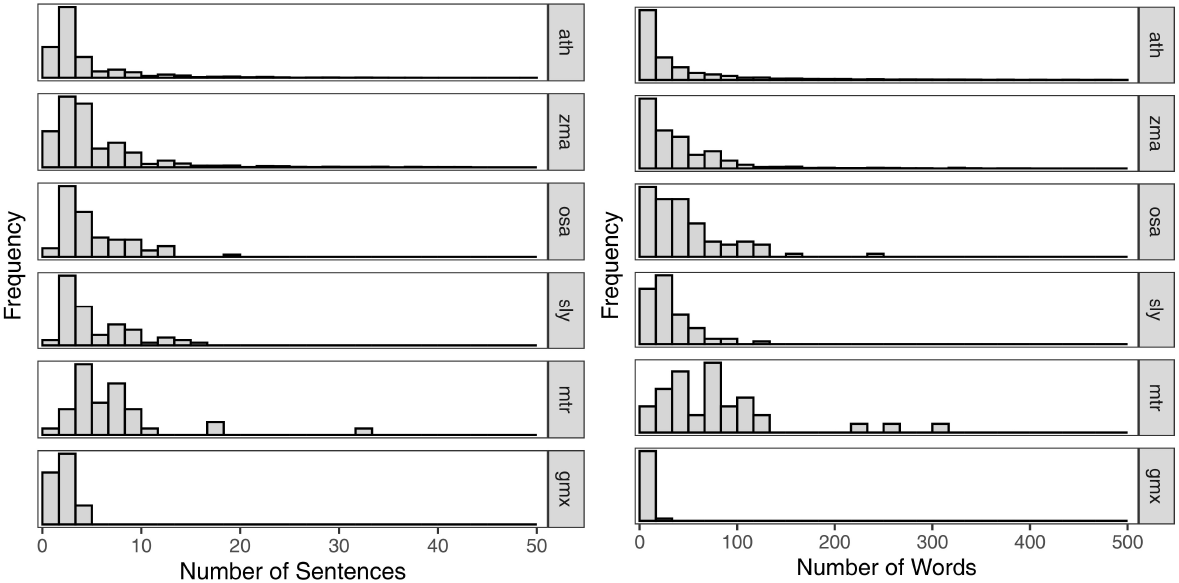
Phenotype description text length distributions across six plant species. The distributions for quantities of text in terms of both sentences (Left) and words (Right) describing phenotypes for genes in each of the plant species. Outliers with very long descriptions are not shown, which includes *<*1% of the genes belonging to Arabidopsis and *<*0.1% of the genes belonging to maize. The y-axis is scaled to be proportional to the quantity of genes for each individual species.

**Figure 2.**
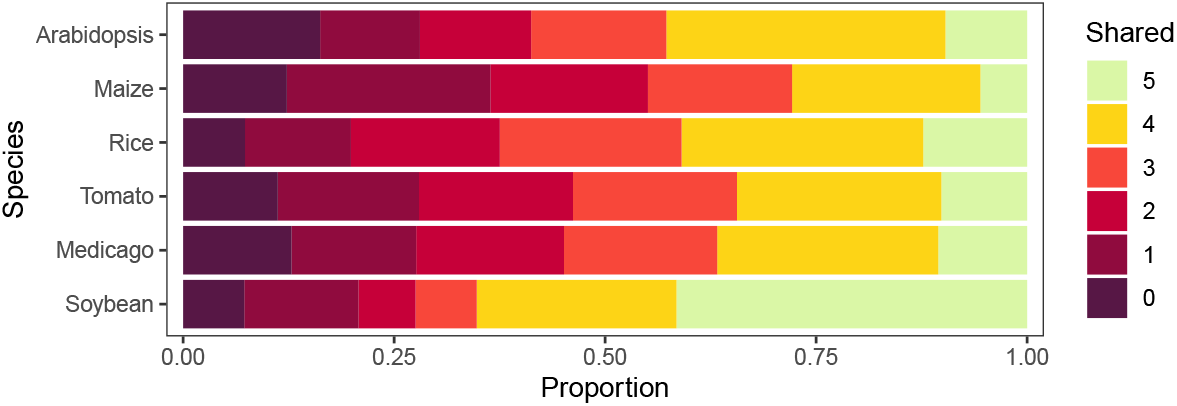
Overlap among vocabularies used to describe phenotypes in each species. For each of the six species in the dataset, listed on the left, the proportion of words in the total vocabulary used in all phenotype descriptions of that species that are shared with the vocabularies of a given additional number of species are shown, with colors indicated on the right. For example, plum/purple indicates the proportion of words used only in that species, and light green indicates the proportion of that species vocabulary that is shared with the vocabulary of all five other species.

For the genes in this dataset, we also collected three types of ontology term annotations: Gene Ontology (GO; Ashburner et al. 2000) annotations, Plant Ontology (PO; Jaiswal et al. 2005; Cooper et al. 2013) annotations, and entity-quality (EQ) statements composed of multiple ontology terms. For in-depth discussion on how EQ statements are composed and compared to one another, see (Hoehndorf et al. 2011; Oellrich et al. 2015; Braun and Lawrence-Dill 2019). GO and PO annotations were additionally sourced from the model species databases (Berardini et al. 2015; Portwood et al. 2019; Wimalanathan et al. 2018) and Planteome (Cooper et al. 2018; http://www.planteome.org), and were limited to those with evidence codes indicating they were either experimentally determined or created through author or curator statements (Consortium 2012; Giglio et al. 2019). The EQ statements were sourced from the dataset of curator-defined EQ statements created by Oellrich et al. 2015. Not all genes in the dataset had at least one annotation of each type, and these quantities are given in Table 1. The preprocessed, merged, and cleaned dataset described here is available and further described through a dedicated repository (see Code and Data Availability).

We also mapped the genes in this dataset to objects from additional bioinformatics resources, namely biochemical pathways in KEGG (Kanehisa et al. 2002) and PlantCyc (Schläpfer et al. 2017), protein-protein associations in STRING (Szklarczyk et al. 2016), ortholog relationships in PANTHER (Thomas et al. 2003), and a hierarchical Arabidopsis gene classification based on phenotypes (Lloyd and Meinke 2012). A subset of the genes in the complete dataset are found in each of these resources (Table 1).

### 2.2 Measure of gene pair similarity

We used a set of approaches for generating *n* by *n* pairwise similarity matrices, where *n* is the number of genes in the dataset, and the values in the matrix are some measure of the similarity between a given pair of genes. Each approach yields one matrix. The approaches belong to two main groups: text-based approaches that translate the text descriptions of phenotype(s) associated with each gene into numerical vectors, so that gene pair similarity can then be found using cosine similarity, and curator-based approaches, that rely on similarities between existing annotations for each gene (GO terms, PO terms, or EQ statements) to quantify gene pair similarity. Each of the text-based approaches used is described in overview here, as well as how the curator-based approaches determine gene pair similarity from annotations.

#### 2.2.1 Tokenizing sentences

For each of the text-based approaches, we determined the effects of treating the entirety of the phenotype descriptions associated with a gene as one concatenated text, and comparing between those texts for pairs of genes to measure gene pair similarity, or by first tokenizing (separating) the phenotype descriptions into individual sentences, and treating those sentences as individual text instances. Then the maximum similarity scores obtained by any pair of sentences was taken as the gene pair relatedness score. This measure is intended to alleviate the effects of genes with longer phenotype descriptions seeming to appear unrelated to ones with shorter ones, and is analogous to looking for local alignments in the text, rather than global ones. In the subsequent Methods sections, we use the word ‘text’, to mean either the concatenation of all phenotype descriptions associated with a gene, or a single sentence from those descriptions, depending on which of these two methods is being described. Sentence tokenization was done with the NLTK package (Bird et al. 2009).

#### 2.2.2 Baseline approach

Some genes in the collected dataset have identical phenotype descriptions. As a baseline approach against which to compare the subsequently described approaches, we include an approach that simply yields a similarity value of 1 for gene pairs that have identical texts, and 0 for gene pairs with texts that differ in any way, after preprocessing.

#### 2.2.3 TF-IDF

Constructing tf-idf (term frequency-inverse document frequency) vectors is one of simplest ways of representing text in a computable format. With this approach, phenotype descriptions are treated as a bag-of-words, and translated to a vector which is the same length as the total number of unique words in the dataset vocabulary, where each position in the vector corresponds to a particular word. The value at the position in the vector for a particular word is the number of times that word appears in the phenotype description (term frequency) weighted by the inverse of the fraction of phenotype descriptions in which that word appears (inverse document frequency). Weighting by the inverse document frequency emphasizes the importance of rarer words (e.g., ‘gametophyte’) and de-emphasizes the importance of more common words (e.g., ‘plant’) in the vector encoding. In addition to this straightforward implementation of the tf-idf approach, we also used as a bigram approach where positions in the vector represent a sequence of two consecutive words (as opposed to the unigram approach, where positions are a single word, as described above). We also used a tf-idf monogram approach where the phenotype descriptions in the datasets are first subset to only include words that are over-represented in journal articles abstracts related to plant phenotypes. The criteria for inclusion was that a word appeared at least twice as frequently in the dataset of plant phenotype related abstracts compared to a general domain corpus. In all cases, cosine similarity was used to calculate gene pair similarity after phenotype descriptions were translated into vectors.

#### 2.2.4 Computational annotation (NOBLE Coder)

NOBLE Coder (Tseytlin et al. 2016) is a computational tool for annotating text with ontology terms. We used NOBLE Coder to annotate phenotype descriptions with terms from a set of bio-ontologies (GO, PO, and PATO), inheriting additional terms using the hierarchical structure of the ontologies. We used NOBLE Coder with both the exact and partial match parameters, which alters how strictly an ontology term must match a text string for an annotation to be assigned. After assigning terms to phenotype descriptions for genes by this method, each gene is represented by a set of terms rather than a set words, and the process of translating these representations into numerical vectors and calculating gene pair relatedness using cosine similarity is the same as with the tf-idf approach, with positions in the resulting vectors referencing terms instead of words. Again, cosine similarity was applied to yield similarity matrices from these resulting vectors.

#### 2.2.5 Topic modeling (LDA and NMF)

We used Latent Dirichlet-Allocation (LDA; Blei et al. 2003) and Non-negative Matrix Factorization (NMF; Lee and Seung 1999) to perform topic modelling on the dataset of phenotype descriptions. These are decomposition algorithms that are widely used in NLP applications (reviewed in Jelodar et al. 2019), and result in translating a document-term matrix into a document-topic matrix (in our case, documents are phenotype descriptions). If the algorithm is run to learn 10 topics, then the outcome is that each phenotype is represented by a vector of length 10 where each position indicates the probability that the phenotype is derived from that particular topic. Determining the appropriate number of topics to use for a particular dataset is often a matter of trying a range of values, and looking at which value produces the most coherent or logical topics given the subject matter. Based on the word probability distributions created using a range of topic quantities, we used our best judgement to elect to use 50 topics and 100 topics for our embedding approaches using each of these algorithms.

#### 2.2.6 Neural network-based embeddings (Word2Vec, Doc2Vec, BERT, BioBERT)

We also used machine learning approaches designed to find vector embeddings that represent the semantics of input text in a compressed space, with positions in the embedding representing abstract semantic features. Word2Vec (Mikolov et al. 2013) is an approach for generating word embeddings based on the contexts in which words appear in a corpus. We used a skip-gram model, where a shallow network is trained to take one word at a time from our corpus as input and predict surrounding context words. The result of this self-supervised training step is a vector embedding for each word that occurs in the dataset of descriptions that reflects the context those words appear in, in a compressed feature space (200 dimensions). To supplement our dataset of phenotype descriptions to build a larger corpus for self-supervised training, we shuffled in sentences accessed from PubMed that were present in abstracts retrieved with queries for the word ‘phenotype’ and any of the names of the species present in our dataset. Hyperparameters for model construction were selected through a validation task of predicting whether ontology term names and synonyms from PATO and PO were parent-child or sibling pairs, or more distantly related. This validation task led to the selection of a skip-gram model using a window size of 8, and a hidden layer size of 200 (see genism package (Rehurek and Sojka 2010) for parameter details). In addition, as a point of comparison, we also used pre-trained published models trained on PubMed (Moen and Ananiadou 2013) and Wikipedia (Lau and Baldwin 2016).

Doc2Vec is an extension of Word2Vec that either exclusively learns embeddings for documents (texts with multiple words) or learns embeddings for documents simultaneously with word embeddings. We used a distributed bag of words architecture where the arbitrary document tags are used as an input in a self-supervised process to predict randomly selected words from the input documents, resulting in network architecture that can be used to infer document-specific embeddings (Le and Mikolov 2014). We utilized the same training approach as for word embeddings, using only concept pairs with multiple words as validation data. In addition, we used a pre-trained Doc2Vec model trained on Wikipedia (Lau and Baldwin 2016).

BERT (Bidirectional Encoder Representations from Transformers) is a large-scale neural network architecture trained on large unlabeled text datasets to predict masked words in sentences and predict whether one sentence follows another in a corpus (Devlin et al. 2018). This results in a network where the encoder can be used to generate context-specific vector embeddings for words in an input sentence. We used both the BERT base model (Devlin et al. 2018) and BioBERT models fine-tuned on abstracts from PubMed and articles from PubMed Central (Lee et al. 2020).

The Doc2Vec models were used to directly infer vector embeddings for phenotype descriptions. The Word2Vec and BERT models generate vector embeddings for each word in phenotype descriptions, so these individual word-embeddings were combined to produce a single vector embedding for each phenotype description. Whether the vectors are summed or averaged is a hyperparameter choice, along with how many encoder layers are used to build the BERT word vectors, and whether those layers should be summed or concatenated. These hyperparameter choices were made using performance on the validation task described previously for the networks trained on phenotype descriptions, and for the pre-trained models we selected hyperparameters based on their performance on a related biomedical sentence similarity problem with the BIOSSES dataset (Soğancıoğlu et al. 2017), and went forward with the hyperparameters that provided the best results on that separate dataset. As with the other approaches, cosine similarity was applied to the resulting vectors to yield similarity matrices.

#### 2.2.7 Using embeddings to generate meaningful vectors with word replacement

Producing the most informative vector representations of phenotype descriptions requires combining the tf-idf approach of explicitly representing the quantity of each particular word from the vocabulary that is present in each phenotype description, and also accounting for semantics through learning vector embeddings of particular words relative to their own meanings in this vocabulary or their meaning relative to the words around them in these phenotypes. We used an approach where pairwise word-similarity matrices for each word in the vocabulary as represented by our Word2Vec models were used to replace each word in all descriptions with the most common word in the vocabulary out of the word itself and the three other most similar words predicted by that model (algorithm detailed in Pontes et al. 2016). This results in substitutions such as ‘susceptible’ to ‘resistance’ that may allow comparisons to be made between phenotypes that simpler bag-of-words approaches would consider as distinct. The resulting vector representations are tf-idf vectors, but the semantic relationships between words as informed by the neural network models is already accounted for prior to encoding.

#### 2.2.8 Curated annotations (GO, PO, EQ statements)

For a point of comparison to the text-based approaches described above, we also used the curator-based annotations to quantify gene pair relatedness. For GO and PO annotations, we calculated similarities as the maximum information content of any single term shared between the annotation sets for a given pair of genes. The more similar two sets of annotations are, the more specific (with higher information content) the terms shared between the two sets will be with respect to the ontology graph structure, leading to greater similarity. In this case, information content is transformed to be in the range of 0 to 1, so that it can be used as a similarity metric compatible with the other approaches used. To quantify similarity between genes using EQ statements, we used the pairwise similarities provided in Oellrich et al. 2015.

### 2.3 Formulating Biologically Relevant Questions

We used additional bioinformatic resources (KEGG, PlantCyc, STRING, PANTHER, etc.) to assess representation of biologically relevant relationships between gene pairs in the dataset, that each approach described above can attempt to recover by quantifying the similarity for that pair of genes, allowing for direct comparison among the approaches (Table 2). Because not all genes in the dataset map to each resource (Table 1), the number of gene pairs that are applicable to each question are not consistent (Table 3). Although these questions are likely related to one another in terms of true biology (e.g., if a pair of genes are related to the same observable phenotype, they are probably more likely to act in a shared pathway), these questions are neither identical nor redundant in the context of this work, because different questions apply to different portions of the dataset, and even within the overlaps of gene pairs that apply to multiple questions, the set of positives (gene pairs for which the correct answer is ‘true’) are not the same (Table 4). For example, the two most similar tasks are ‘Associations’ and ‘Pathways’, where 1,271,297 of the same gene pairs are considered in both tasks, and the Jaccard similarity between the two sets of target values (‘true’, ‘false’) between those gene pairs is only 0.172 (Table 4). For this reason, we looked at the results of each of these questions individually rather than combining them.

**Table 2.** Biological relationships tested in each task.

| Task | Description (Are genes A and B...) | Knowledge Source |
| --- | --- | --- |
| Phenotypes | ...impacting the same phenotype? | Lloyd and Meinke, 2012 |
| Pathways | ...functioning in the same pathway? | PlantCyc, KEGG |
| Associations | ...known to share a function or process? | STRING |
| Orthologs | ...orthologous to one another? | PANTHER |

**Table 3.** Number of genes and gene pairs used for each task.

| Question | All Text Data |  |  |  | With Annotations |  |  |  |
| --- | --- | --- | --- | --- | --- | --- | --- | --- |
|  | Genes | Pairs | Positive Pairs |  | Genes | Pairs | Positive Pairs |  |
| Phenotypes | 2,356 | 2,774,190 | 303,009 | 10.92% | 2,284 | 2,607,186 | 293,221 | 11.25% |
| Pathways | 1,838 | 1,688,203 | 45,847 | 2.72% | 1,045 | 545,490 | 14,853 | 2.72% |
| Associations | 4,620 | 9,343,325 | 147,271 | 1.58% | 2,377 | 2,530,556 | 52,541 | 2.08% |
| Orthologs | 921 | 248,913 | 65 | 0.03% | 368 | 43,187 | 23 | 0.05% |

**Table 4.** Similarities among datasets across biological tasks.

| Task 1 | Task 2 | Overlap Size | Jaccard (Pairs) | Jaccard (Values) |
| --- | --- | --- | --- | --- |
| Associations | Pathways | 1,271,297 | 0.130 | 0.172 |
| Phenotypes | Pathways | 511,566 | 0.129 | 0.050 |
| Phenotypes | Associations | 2,687,721 | 0.285 | 0.032 |
| Pathways | Orthologs | 29,654 | 0.016 | 0.012 |
| Phenotypes | Orthologs | 0 | 0.000 |  |
| Associations | Orthologs | 0 | 0.000 |  |

## 3 RESULTS

### 3.1 Text-based approaches recover biological relationships

Using each of the text-based approaches as well as using similarity metrics over the existing curated annotations, we calculated gene pair similarity values for all pairs of genes in our dataset. We measured the success of each approach for (1) predicting whether two genes were orthologs (as specified in PANTHER), (2) predicting known protein associations specified in STRING, (3) predicting whether two genes functioned in at least one of the same biochemical pathways (as specified in PlantCyc and KEGG), and (4) at predicting whether two Arabidopsis genes belonged to one of the phenotype categories specified by Lloyd and Meinke 2012. For each of these biological questions, a given approach for measuring gene similarity is considered useful if the distribution of values for gene pairs for which the answer to the question is true is distinct from the distribution of values for gene pairs for which the answer to question is false. The success of each approach for each biological question was calculated in terms of the maximum F_1_ statistic. We also recalculated the maximum F_1_ statistic for just the genes for which we have GO annotations, PO annotations, and EQ statements, to directly compare performance of each approach on each question with approaches that are based on curation (Table 5, Supplemental Table S1).

**Table 5.** Comparing F_1_scores and group significance rates for phenotype and pathway relationships.

| Approach | Category | Concat | Phenotypes ($F_1$, % Significant) | | | | Pathways ($F_1$, % Significant) | | | |
| --- | --- | --- | --- | --- | --- | --- | --- | --- | --- | --- |
|  |  |  | All Genes |  | Curated |  | All Genes |  | Curated |  |
| Baseline | Baseline | Yes | 0.197 | 17% | 0.202 | 19% | 0.052 | 7% | 0.053 | 3% |
| TF-IDF (Unigrams) | TF-IDF | Yes | 0.465 | 100% | 0.473 | 100% | 0.105 | 61% | 0.114 | 57% |
| TF-IDF (Unigrams & Bigrams) | TF-IDF | Yes | 0.473 | 100% | 0.482 | 100% | 0.109 | 64% | 0.120 | 58% |
| TF-IDF (Plant Article Unigrams) | TF-IDF | Yes | 0.462 | 100% | 0.471 | 100% | 0.100 | 59% | 0.110 | 59% |
| NOBLE Coder (Precise) | Annotation | Yes | 0.364 | 91% | 0.370 | 91% | 0.075 | 34% | 0.079 | 23% |
| NOBLE Coder (Partial) | Annotation | Yes | 0.372 | 100% | 0.380 | 100% | 0.087 | 42% | 0.094 | 33% |
| LDA (50 Topics) | Topic Modeling | Yes | 0.327 | 88% | 0.336 | 88% | 0.075 | 12% | 0.088 | 24% |
| LDA (100 Topics) | Topic Modeling | Yes | 0.329 | 88% | 0.337 | 91% | 0.072 | 25% | 0.083 | 26% |
| NMF (50 Topics) | Topic Modeling | Yes | 0.436 | 100% | 0.448 | 100% | 0.090 | 44% | 0.103 | 46% |
| NMF (100 Topics) | Topic Modeling | Yes | 0.418 | 100% | 0.423 | 100% | 0.093 | 49% | 0.102 | 48% |
| Doc2Vec (Wikipedia) | ML (Embeddings) | Yes | 0.313 | 86% | 0.322 | 86% | 0.068 | 22% | 0.068 | 15% |
| Doc2Vec (Plants) | ML (Embeddings) | Yes | 0.237 | 45% | 0.240 | 50% | 0.059 | 20% | 0.060 | 15% |
| Word2Vec (Wikipedia) | ML (Embeddings) | Yes | 0.276 | 98% | 0.284 | 98% | 0.087 | 12% | 0.096 | 21% |
| Word2Vec (PubMed) | ML (Embeddings) | Yes | 0.320 | 93% | 0.327 | 98% | 0.093 | 14% | 0.108 | 21% |
| Word2Vec (Plants) | ML (Embeddings) | Yes | 0.443 | 98% | 0.452 | 98% | 0.102 | 26% | 0.111 | 36% |
| BERT | ML (Embeddings) | Yes | 0.289 | 93% | 0.296 | 95% | 0.084 | 4% | 0.096 | 15% |
| BioBERT | ML (Embeddings) | Yes | 0.309 | 81% | 0.316 | 83% | 0.087 | 5% | 0.098 | 17% |
| Word2Vec (Wikipedia) | ML (Word Replacement) | Yes | 0.387 | 98% | 0.396 | 100% | 0.099 | 46% | 0.107 | 43% |
| Word2Vec (PubMed) | ML (Word Replacement) | Yes | 0.417 | 100% | 0.426 | 100% | 0.104 | 50% | 0.111 | 45% |
| Word2Vec (Plant Phenotypes) | ML (Word Replacement) | Yes | 0.456 | 98% | 0.464 | 100% | 0.103 | 59% | 0.114 | 53% |
| Baseline | Baseline | No | 0.465 | 74% | 0.476 | 76% | 0.089 | 19% | 0.097 | 39% |
| TF-IDF (Unigrams) | TF-IDF | No | 0.544 | 95% | 0.555 | 100% | 0.100 | 51% | 0.108 | 61% |
| TF-IDF (Unigrams & Bigrams) | TF-IDF | No | 0.540 | 95% | 0.552 | 95% | 0.100 | 58% | 0.107 | 63% |
| TF-IDF (Plant Article Unigrams) | TF-IDF | No | 0.555 | 98% | 0.562 | 100% | 0.098 | 33% | 0.106 | 52% |
| NOBLE Coder (Precise) | Annotation | No | 0.457 | 83% | 0.466 | 83% | 0.089 | 5% | 0.103 | 33% |
| NOBLE Coder (Partial) | Annotation | No | 0.501 | 95% | 0.510 | 98% | 0.095 | 24% | 0.102 | 47% |
| NMF (50 Topics) | Topic Modeling | No | 0.509 | 88% | 0.520 | 88% | 0.092 | 13% | 0.099 | 30% |
| NMF (100 Topics) | Topic Modeling | No | 0.517 | 93% | 0.527 | 95% | 0.090 | 18% | 0.100 | 37% |
| LDA (50 Topics) | Topic Modeling | No | 0.500 | 98% | 0.511 | 100% | 0.089 | 11% | 0.099 | 35% |
| LDA (100 Topics) | Topic Modeling | No | 0.500 | 98% | 0.510 | 100% | 0.097 | 19% | 0.104 | 41% |
| Doc2Vec (Wikipedia) | ML (Embeddings) | No | 0.518 | 100% | 0.529 | 100% | 0.100 | 23% | 0.107 | 51% |
| Doc2Vec (Plants) | ML (Embeddings) | No | 0.561 | 95% | 0.571 | 95% | 0.098 | 26% | 0.104 | 52% |
| Word2Vec (Wikipedia) | ML (Embeddings) | No | 0.521 | 98% | 0.532 | 98% | 0.098 | 12% | 0.106 | 38% |
| Word2Vec (PubMed) | ML (Embeddings) | No | 0.529 | 95% | 0.540 | 100% | 0.100 | 19% | 0.108 | 42% |
| Word2Vec (Plants) | ML (Embeddings) | No | 0.558 | 98% | 0.569 | 98% | 0.103 | 30% | 0.111 | 51% |
| BERT | ML (Embeddings) | No | 0.500 | 100% | 0.511 | 100% | 0.102 | 25% | 0.111 | 43% |
| BioBERT | ML (Embeddings) | No | 0.515 | 100% | 0.526 | 100% | 0.104 | 24% | 0.113 | 48% |
| Word2Vec (Wikipedia) | ML (Word Replacement) | No | 0.558 | 100% | 0.568 | 100% | 0.102 | 33% | 0.110 | 53% |
| Word2Vec (PubMed) | ML (Word Replacement) | No | 0.554 | 100% | 0.566 | 100% | 0.101 | 34% | 0.109 | 51% |
| Word2Vec (Plant Phenotypes) | ML (Word Replacement) | No | 0.566 | 95% | 0.577 | 93% | 0.099 | 37% | 0.107 | 51% |
| GO | Curation |  |  |  | 0.249 | 64% |  |  | 0.140 | 38% |
| PO | Curation |  |  |  | 0.215 | 17% |  |  | 0.056 | 9% |
| EQs | Curation |  |  |  | 0.475 | 74% |  |  | 0.093 | 51% |

#### 3.1.1 Text-based approach performance is dependent on biological query type

Of the four biological questions assessed for this analysis, predicting whether two genes were orthologous, whether two genes shared an association, or whether two genes belonged to a shared biochemical pathway were infeasible for any of the text-based or curation-based approaches, in terms of broad performance measured with maximum F_1_ statistics (Table 5, Supplemental Table S1). The largest F_1_ statistic obtained across all three of these tasks for any approach was 0.140 using the curated GO annotations, with all other approaches yielding F_1_ values less than 0.12 (Table 5, Supplemental Table S1). However, F_1_ statistics were much higher for the task of predicting whether two genes belonged to the same phenotypic category, an expected result given that this prediction follows directly from the explicit contents of the phenotypic descriptions (Table 5). This was true for both the text-based and curation-based approaches, but the best performance was achieved using text-based approaches (Table 5). Performance on this task of predicting whether two genes share a phenotypic category can be broken down by general classes of approach (Figure 3).

**Figure 3.**
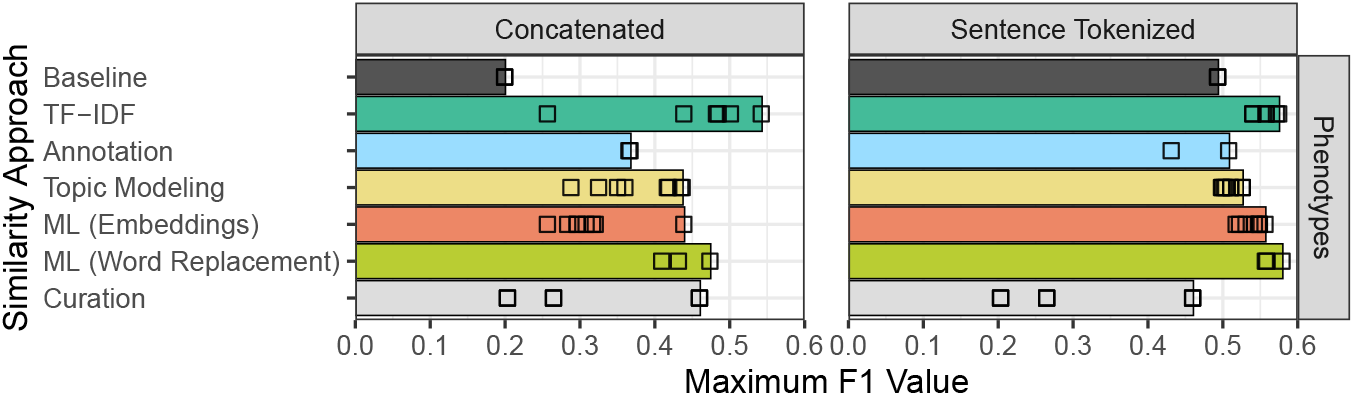
Comparing the groups of gene pair similarity approaches. The maximum F_1_ statistics for each approach in each broad category for measuring gene similarity is shown, with the bar indicating the best F_1_ statistics among all the approaches in that general group. Bars on the left indicate performance when phenotype descriptions are treated as one concatenated piece of text, and bars on the right indicate performance when the descriptions are sentence tokenized first.

As previously stated, all approaches were unsuccessful in predicting ortholog relationships (Supplemental Table S1). In addition, all approaches were completely unsuccessful in predicting whether two genes from different species were involved in a common biochemical pathway (Supplemental Table S2). Even though the maximum F_1_ statistics for predicting whether two genes share a pathway were already low, these values were even lower when filtering the dataset to only look at interspecies gene pairs, and marginally greater when filtering the dataset to only look at intraspecies gene pairs (Supplemental Table S2). Therefore, even the very small amount of biological information recovered only applies to looking at genes from within the same species. This indicates that comparing the text of phenotype descriptions across different species is not biologically informative in this case. This might not be true for all species or all phenotypes, but it does not generalize across the current dataset of available plant phenotype descriptions.

#### 3.1.2 Significant description similarity within individual phenotype and pathway gene groups

We evaluated similarities of phenotypic descriptions as a percentile based on F_1_ score, and to visualize the results, plotted the average gene-to-gene similarity for phenotypic categories (Figure 4; top panel). Next, we imposed the same evaluation and visualization for genes mapped to each individual pathway (Figure 4; bottom two panels). Although predicting whether two genes shared a biochemical pathway was generally unsuccessful (low maximum F_1_), this is in part a consequence of the fact that pathways vary greatly in how related the phenotype descriptions for their component genes are. In Table 5 we report calculated p-values. This was accomplished by randomly sampling groups of genes at each value of *n* then calculating p-values for each phenotype category and pathway based on the probability of each approach generating a mean similarity value between genes in that group that is that large or larger, controlling for false discovery rate for each approach with the Benjamini–Hochberg procedure. For text-based approaches using sentence tokenization, 81% to 100% of the phenotypic categories had a significantly large average similarity value (with respect to the Benjamini–Hochberg procedure), while between 6% and 39% of the pathways obtained significant average similarity values, for these same approaches, with an average of 23% (Table 5). Taken together, these results indicate that while text-based similarity values are not broadly indicative of whether or not two genes share a pathway, there is a significant subset of known pathways for which this is the case. In the case of groups of genes belonging to the same pathway that do have similar phenotype descriptions, these are generally either due to mentions of downstream phenotypic effects of pathway disruption, or more direct mentions of the pathway function or role. For example, the descriptions associated with genes in the chlorophyll degradation pathway include mentions of necrotic lesions, and the descriptions associated with genes in the phospholipid desaturation pathway include mentions of fatty acid levels or composition.

**Figure 4.**
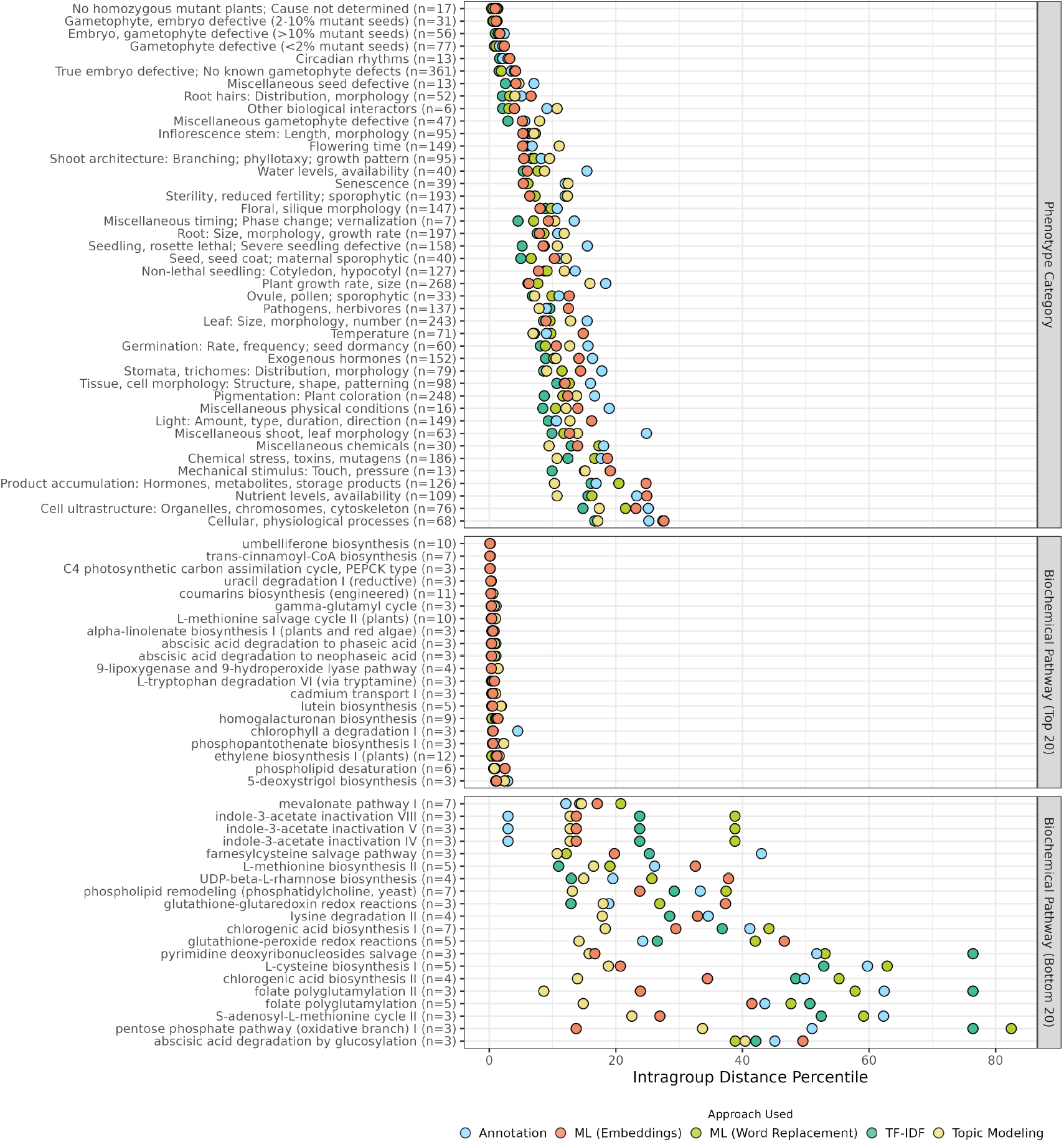
Cohesiveness of phenotype and pathway gene groups. Phenotype categories (Top) and Biochemical Pathways (Middle and Bottom) are listed, with the number of genes in these datasets belonging to each group listed to the right of the group’s name. The x-axis indicates group cohesiveness, given as the percentile against all pairwise gene distances that the average distance between any two genes in that group falls in. The minimum value of this metric achieved by any approach that is in the listed category is shown. For example, the location of the yellow dot in a particular row indicates the smallest intragroup distance percentile obtained by any approach in the topic modeling category of text-based approaches for that particular group of genes.

#### 3.1.3 Combining syntactic and semantic approaches improves recovery of phenotypic categories

The purely syntactic text-based approaches (tf-idf) were among the most successful in terms of maximum F_1_ statistic for predicting whether gene pairs belonged to the same phenotypic category (Table 5, Figure 3). In general, semantic approaches that use ML techniques to drastically reduce the dimensionality of the vector encoding for each text instance were comparably successful (Table 5, Figure 3). However, the combined approaches where semantic techniques were used to augment the information in the tf-idf vectors by replacing words with similar words prior to encoding provided a boost in performance over other approaches (Table 5, Figure 3). Taken together, this indicates that this dataset contains phenotype descriptions for genes in the same phenotypic category that are similar both in terms of explicitly shared words (where syntactic approaches are most helpful), as well as genes that are similar only in terms of shared meaning but not specific words (where semantic approaches provide an advantage). Using word embedding models trained on plant phenotype specific data provided marginal improvement over models trained on PubMed generally or the Wikipedia corpus, but all three models provided the same boost over other approaches when applied to word replacement, indicating that useful associations between words for recovering common phenotypic categories from descriptions are not limited to relationships only represented in a narrow corpus of text related to plant phenotypes. Given that using bio-ontologies for this same task did not perform as well as text-based approaches, and one of the main functions of such ontologies in this case is to inject domain-specific inferences into the similarity metrics, this result is not surprising.

#### 3.1.4 Sentence tokenization is important for comparing phenotypes

For all the text-based approaches on all the biological questions posed, the preprocessing step of tokenizing phenotype descriptions into sentences and evaluating gene pair relatedness as the maximum pairwise sentence similarity resulted in greater F_1_ statistics (Table 5, Supplemental Table S1). Unexpectedly, this held true even for approaches that are generally intended for use with larger input texts, such as Doc2Vec, and topic modeling algorithms LDA and NMF. This indicates that when predicting whether two genes share a common role, it is important to account for ‘local alignments’ in their associated phenotype descriptions, as the similarity might exist between single sentences associated with those genes while other sentences act as noise obscuring this relationship.

### 3.2 Enabling biologists to use these methods and dataset

#### 3.2.1 Web application (QuOATS)

We have developed a web application called QuOATS (Querying with Ontology Annotations and Text Similarity) for querying the dataset described here through leveraging the computational methods described here (Figure 5A). The underlying dataset of plant genes is the same as is described previously (Table 1), and can be filtered to include particular species (Figure 5B). The application supports four different query types (Figure 5D), with the primary purpose being to obtain lists of genes that are related to phenotypes described similarly to some phenotypic characteristic(s) of interest. Firstly, a free text query can be used to search the dataset for any genes related to phenotypes that are described similarly to text strings separated by periods in the query (Figure 5E). Secondly, a keyword query can be used to input any number of strings of any length, and genes whose phenotype descriptions contain those strings (after preprocessing including stemming and case-normalization) are returned (Figure 5F). Thirdly, an ontology term query can be used to search for any genes annotated by curators with one or more ontology terms, either directly or inherited through the ontology hierarchy (Figure 5G). Lastly, a gene identifier query can be carried out to search for any gene name, protein name, gene model, or any other gene identifier potentially represented in the dataset. Selecting a gene from the returned list of candidates that match the query will auto-complete a second query that returns genes related to phenotypes that are described similarly to the selected genes (Figure 5H).The similarity scores used to rank genes in the returned list are calculated using approaches described here, selected from a drop-down menu in the web application (Figure 5C).

**Figure 5.**
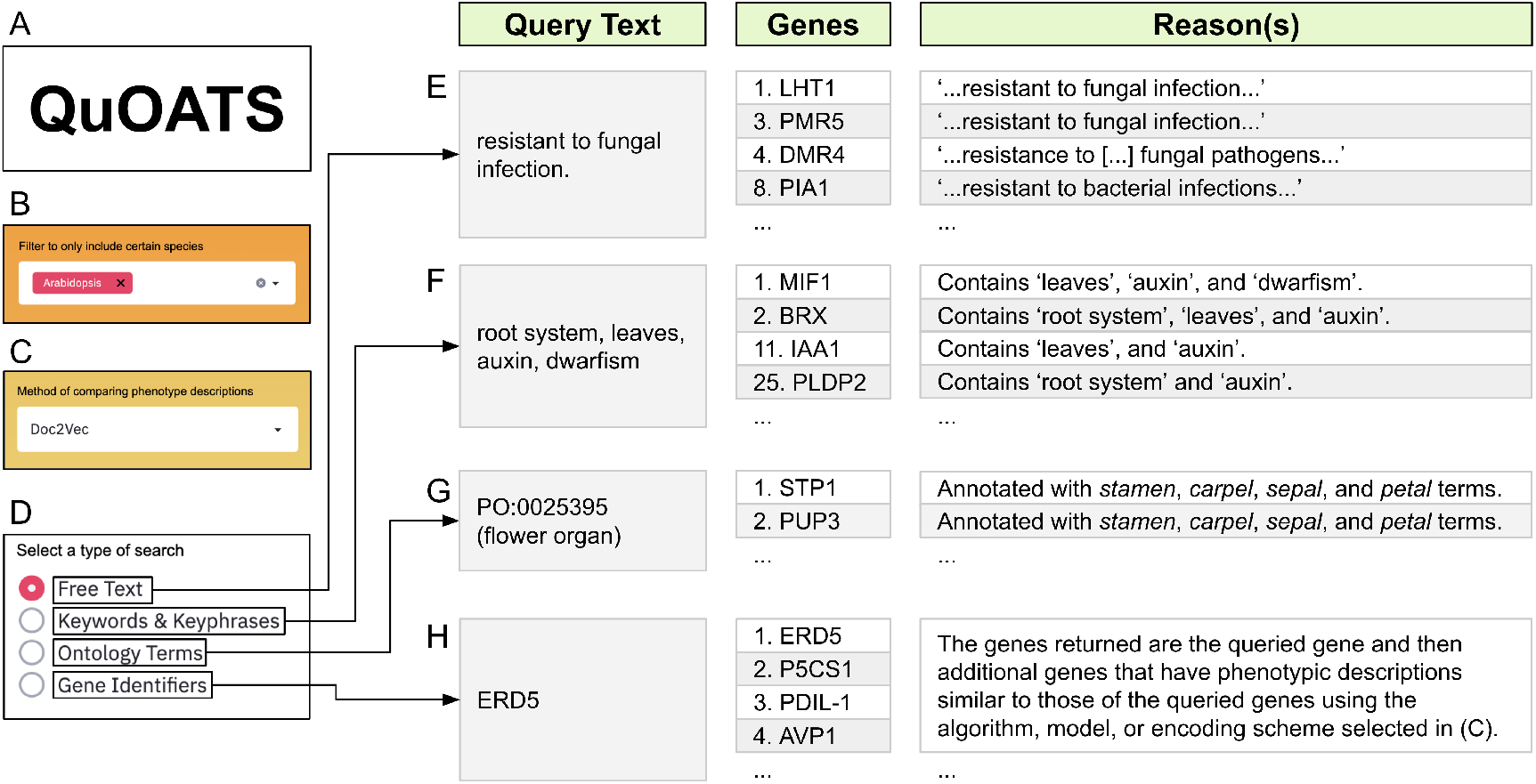
Querying plant genes, annotations, and phenotype descriptions. A. The name of the web application we have developed. B. Option to subset the available dataset to only include certain species. C. Option to select the algorithm or method used to compare phenotype descriptions. D. Four different types of querying are supported. E, F, G, H. The information given here for each query type is presented when using the webtool, but has been re-organized and truncated for the sake of illustration. The queries listed are the text strings that are entered into the search bar to generate the results shown. The returned genes appear in the results in the row indicated by the number to the left of the gene names. The reasons that these genes appear in this order given these particular queries are described to the right of the gene names.

#### 3.2.2 Proof of concept applications of the web tool

In our previous findings illustrated in Braun and Lawrence-Dill 2019, we discussed how a set of genes related to anthocyanin biosynthesis could be used to demonstrate recovering gene groups by querying specifically with phenotype descriptions or computationally generated annotations from those descriptions. Specifically, we looked at a dataset of 16 maize genes (Li et al. 2019) and 21 genes from Arabidopsis (Appelhagen et al. 2014) but only 10 of the maize genes and 16 of the Arabidopsis genes were present in the dataset. Our expanded dataset in this work includes 14 of those maize genes and 18 of the Arabidopsis genes. We now evaluate the results of querying with each of these genes in the web application QuOATS, to recover both genes in the same species from these sets and genes in the alternate species. Over the 64 total queries (32 within the same species and 32 between species), we quantified the average and standard deviation of the number of target genes contained in bins of ranks in the query results, in bin sizes of 10 up to 50, and a final bin for genes that obtain ranks higher than 50 (Figure 6). Additionally, we also repeated this analysis for a set of 9 core autophagy genes in Arabidopsis (Figure 6). These queries illustrate a proof-of-concept whereby the web application can be used to query with phenotypic descriptions associated with one gene to recover other related genes. This application demonstrates the utility of applying text-based algorithms in cases where ontology annotations are either not present, are insufficient, or could simply be augmented by allowing additional, less rigidly-defined phenotype descriptions to be searchable as well.

**Figure 6.**
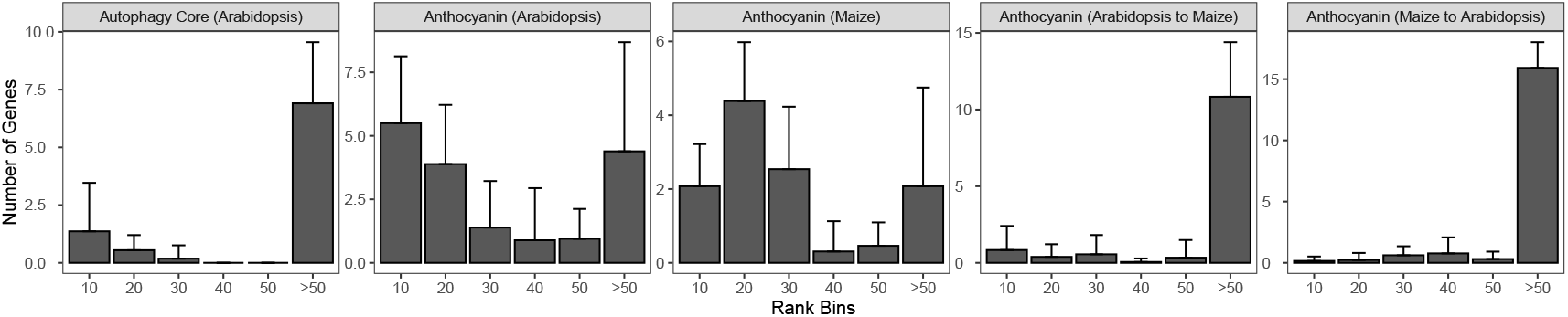
Querying with autophagy core genes and anthocyanin biosynthesis genes in QuOATS. The labels above each plot indicate the set of genes, the species of the genes used as the queries, and then the species for which the resulting ranked genes were filtered (in the case of the left three plots the species is the same for queries and targets). Bars represent bins of rank values for returned genes. Their height indicates the average number of genes with those ranks returned in each query. The error bar indicates the standard deviation in each case. Bars in each plot are labeled with the rank that falls in the right-most edge of that bin. For example, the bar labelled 20 represents genes that were ranked between 11 and 20 in the query results.

## 4 DISCUSSION

The difficulty in computing on phenotypic data is largely a consequence of extreme variability with which these data are represented, and the diversity of ways that phenotypes are measured, quantified, and described. This is in contrast with sequence data; biology as a field has enormously benefited from the ways in which sequence data are intuitively computed on, given the naturally occurring nucleic acid and amino acid coding systems. Sequencing technology provided the datasets to compute on, and algorithms and applications like BLAST provided the means to make use of these data (Altschul et al. 1990). Ontologies have begun to provide a similar means for making phenotypic data computable, and processing of natural language provides an additional avenue by which we can make biological inferences if we have the datasets on which to apply them. The combination of biological ontologies, machine learning approaches, and NLP provide strategies for handling phenotypic descriptions and learning from it where it exists.

Plant phenotypes are frequently described as text within academic papers or research notes. However, these text descriptions are rarely incorporated into relevant research community databases, associated with a specific gene or genotype, and made readily available as part of the growing data resources for that species. This could be the case for a variety of reasons, including the difficulties involved with extracting phenotype descriptions from larger texts, the curatorial effort necessary to produce high quality datasets of phenotypes descriptions associated with genes, or because these text representations of phenotypes are considered a non-valuable data type, and are instead represented by annotations using structured vocabularies of hierarchical terms such as biological ontologies. Notable exceptions to this situation exist, including The Arabidopsis Information Resource (TAIR), which contains thousands of text descriptions of phenotypes mapped to specific Arabidopsis genes (Berardini et al. 2015).

In this work, we have shown that a variety of NLP approaches for vectorizing phenotype descriptions in order to generate gene pair similarity matrices are equally or more predictive in general of known phenotype categorizations compared to using existing curated annotations for this task. Based on these results, we argue that it is worthwhile for databases that contain gene-to-phenotype information to include natural language descriptions of phenotypes. In addition, these descriptions should be made accessible to emergent AI tools, like ChatGPT, to enable the creation of additional resources that can make text descriptions of plant phenotypes more findable and accessible for biological analytics.

The natural language descriptions of phenotypes are useful, and when combined with NLP approaches for computationally representing text can be leveraged to provide a way for researchers to quickly identify genes associated with phenotypes similar to the ones that they are observing or studying. Natural language descriptions can also be used to organize genes computationally on a large scale and discover which categorizations of phenotypes are present in a dataset, with techniques like clustering and topic modeling. In some cases, this natural language data may be easier to generate than ontology annotations. In situations where curators are not available (or have limited time) to generate the high-confidence ontology term annotation datasets, it may be faster or still possible for authors or someone else to at least identify the free-text portions of the manuscript that include phenotype descriptions, and the genes associated with them. In the near future, NLP techniques for parsing full-texts may also progress to the point where this phenotype identification could be done automatically as well. In these instances, we argue it is worthwhile to generate and make accessible this free text phenotype information. In other cases, these text data might already be generated, but are potentially discarded. In situations where curators are actively involved in generating ontology annotations from papers, this process often involves the tasks of highlighting text from the paper, or possibly writing down the phenotype descriptions first then producing the ontology term representation of those associations. Given that the free text itself is useful, we argue it should be retained in the final mapping in the resulting database or dataset rather than being discarded as an intermediate data form. It is possible that for some applications the ontology annotations will be more useful than the natural language descriptions, for example when making comparisons between species, but we have shown that this is not always the case, and if it is being generated regardless, it makes sense to retain the natural language and make it available.

The area where the application of these methods would likely make the most difference is for species where phenotypes are still largely described in general biological terms rather than in cases where phenotype descriptions are limited to special vocabularies and/or where phenotype data have been carefully curated in a pervasive, large-scale way (as is the case with human phenotypes and diseases). In addition, these approaches would make the most sense to use when high quality, curated datasets of ontologized phenotypes are either not available or curation into those data forms are not financially feasible. In these cases, if at a minimum phenotype descriptions are extracted from literature and associated with specific genes in an accessible community database, NLP methods can be applied to organize these data and to group by genes into sets that impact similar phenotypes, therefore allowing researchers to search based on linguistic similarity.

Not only do plant scientists understand their phenotypes and use rich language to describe them, there is a diversity of algorithms available to enable computation on phenotypic descriptions so that the scope of data any single researcher can access becomes quite expansive. In 2011, Mike Freeling made an impression by saying, “Ontologies are for people who don’t understand their phenotype,” to CJLD at the Annual Maize Meeting Genetics Conference in response to a request to review the completeness of the MaizeGDB Phenotypic Controlled Vocabulary (Michael Freeling personal communication). While ontologies have proved invaluable for managing and analyzing the massive quantity of data that biologists deal with, we think that this quote emphasizes the key finding for the efforts here: that we should not undervalue the utility of free text as a datatype, and that it should be made available through bioinformatic resources that provide phenotypic data to the research community, given that we have the computational tools to leverage it in useful ways.

## DATA AND CODE AVAILABILITY

The dataset of plant genes collected from other sources for this work is available at https://github.com/Dill-PICL/Plant-data, along with all the code for preprocessing, reshaping, and merging this data. The code for carrying out the analysis shown here has its own repository at https://github.com/Dill-PICL/NLP-Plant-Phenotypes. The results given here can be reproduced using code and datasets at those locations. In addition, a Python package called OATS (Ontology Annotation and Text Similarity) for working with gene-phenotype datasets, ontology annotations, and free-text was developed in parallel with this work. This package was used extensively for this analysis, and can be found at https://git.io/JTuqv, with documentation available at https://irbraun-oats.readthedocs.io. We have combined the dataset and some of the techniques for identifying similar texts into a streamlit web application named QuOATS available at https://quoats.dill-picl.org/. Use this tool for looking up genes by phenotype keywords or phrases, or finding genes with similar descriptions to a searched phenotype description. The code for this web application is available at https://git.io/Jtv9J.

## AUTHOR CONTRIBUTION

IRB and CJLD conceived the idea for this work. IRB created the first demonstration of all results and drafted the initial version of the manuscript. DCB conceived and reviewed autophagy analyses included as a proof of concept. CFY directed the reproducibility review and edited the manuscript, accordingly. JPDR reproduced previous results and updated the manuscript accordingly. All authors read, edited, and approved the manuscript.

## ACKNOWLEDGEMENTS

The authors thank Carson Andorf, John Portwood, and Naama Menda for their assistance in obtaining the dataset of plant phenotype descriptions and annotations. The authors thank Darwin Campbell and Scott Zarecor for their assistance with dataset creation and hosting and maintenance of the web application discussed here. The authors thank Marna Yandeau-Nelson, Iddo Friedberg, Baskar Ganapathysubramanian, Annette O’Connor, and Qi Li for useful discussions on the topics discussed here, and particularly thank Marna Yandeau-Nelson for assistance in editing and Iddo Friedberg for assistance in reviewing the design of the web application. The authors thank Leila Fattel for helpful discussions and support. The authors also thank the High Performance Computing support team at Iowa State University for their help throughout the project.

## REFERENCES

Altschul, S. F., Gish, W., Miller, W., Myers, E. W., and Lipman, D. J. (1990). Basic local alignment search tool. Journal of Molecular Biology, 215(3):403–410.

Appelhagen, I., Thiedig, K., Nordholt, N., Schmidt, N., Huep, G., Sagasser, M., and Weisshaar, B. (2014). Update on transparent testa mutants from arabidopsis thaliana: characterisation of new alleles from an isogenic collection. Planta, 240(5):955–970.

Ashburner, M., Ball, C. A., Blake, J. A., Botstein, D., Butler, H., Cherry, J. M., Davis, A. P., Dolinski, K., Dwight, S. S., Eppig, J. T., et al. (2000). Gene ontology: tool for the unification of biology. Nature genetics, 25(1):25–29.

Berardini, T. Z., Reiser, L., Li, D., Mezheritsky, Y., Muller, R., Strait, E., and Huala, E. (2015). The arabidopsis information resource: making and mining the “gold standard” annotated reference plant genome. genesis, 53(8):474–485.

Bird, S., Klein, E., and Loper, E. (2009). Natural language processing with Python: analyzing text with the natural language toolkit. “ O’Reilly Media, Inc.”.

Blei, D. M., Ng, A. Y., and Jordan, M. I. (2003). Latent dirichlet allocation. Journal of machine Learning research, 3(Jan):993–1022.

Braun, I., Balhoff, J. P., Berardini, T. Z., Cooper, L., Gkoutos, G. V., Harper, L. C., Huala, E., Jaiswal, P., Kazic, T., Lapp, H., et al. (2018). ‘computable’phenotypes enable comparative and predictive phenomics among plant species and across domains of life.

Braun, I. R. and Lawrence-Dill, C. J. (2019). Automated methods enable direct computation on phenotypic descriptions for novel candidate gene prediction. Frontiers in Plant Science, 10:1629.

Chen, Q., Peng, Y., and Lu, Z. (2019). Biosentvec: creating sentence embeddings for biomedical texts. In 2019 IEEE International Conference on Healthcare Informatics (ICHI), pages 1–5. IEEE.

Consortium, G. O. (2012). Gene ontology annotations and resources. Nucleic acids research, 41(D1):D530–D535.

Cooper, L., Meier, A., Laporte, M.-A., Elser, J. L., Mungall, C., Sinn, B. T., Cavaliere, D., Carbon, S., Dunn, N. A., Smith, B., et al. (2018). The planteome database: an integrated resource for reference ontologies, plant genomics and phenomics. Nucleic acids research, 46(D1):D1168–D1180.

Cooper, L., Walls, R. L., Elser, J., Gandolfo, M. A., Stevenson, D. W., Smith, B., Preece, J., Athreya, B., Mungall, C. J., Rensing, S., et al. (2013). The plant ontology as a tool for comparative plant anatomy and genomic analyses. Plant and Cell Physiology, 54(2):e1–e1.

De Boom, C., Van Canneyt, S., Demeester, T., and Dhoedt, B. (2016). Representation learning for very short texts using weighted word embedding aggregation. Pattern Recognition Letters, 80:150–156.

Devlin, J., Chang, M.-W., Lee, K., and Toutanova, K. (2018). Bert: Pre-training of deep bidirectional transformers for language understanding. arXiv preprint arXiv:1810.04805.

Dwivedi, Y. K., Kshetri, N., Hughes, L., Slade, E. L., Jeyaraj, A., Kar, A. K., Baabdullah, A. M., Koohang, A., Raghavan, V., Ahuja, M., et al. (2023). “So what if ChatGPT wrote it?” Multidisciplinary perspectives on opportunities, challenges and implications of generative conversational AI for research, practice and policy. International Journal of Information Management, (71):102642.

Edmunds, R. C., Su, B., Balhoff, J. P., Eames, B. F., Dahdul, W. M., Lapp, H., Lundberg, J. G., Vision, T. J., Dunham, R. A., Mabee, P. M., et al. (2015). Phenoscape: identifying candidate genes for evolutionary phenotypes. Molecular biology and evolution, 33(1):13–24.

Fernandez-Pozo, N., Menda, N., Edwards, J. D., Saha, S., Tecle, I. Y., Strickler, S. R., Bombarely, A., Fisher-York, T., Pujar, A., Foerster, H., et al. (2015). The sol genomics network (sgn)—from genotype to phenotype to breeding. Nucleic acids research, 43(D1):D1036–D1041.

Giglio, M., Tauber, R., Nadendla, S., Munro, J., Olley, D., Ball, S., Mitraka, E., Schriml, L. M., Gaudet, P., Hobbs, E. T., et al. (2019). Eco, the evidence & conclusion ontology: community standard for evidence information. Nucleic acids research, 47(D1):D1186–D1194.

Hoehndorf, R., Schofield, P. N., and Gkoutos, G. V. (2011). Phenomenet: a whole-phenome approach to disease gene discovery. Nucleic acids research, 39(18):e119–e119.

Huang, D. W., Sherman, B. T., and Lempicki, R. A. (2009). Bioinformatics enrichment tools: paths toward the comprehensive functional analysis of large gene lists. Nucleic acids research, 37(1):1–13.

Jaiswal, P., Avraham, S., Ilic, K., Kellogg, E. A., McCouch, S., Pujar, A., Reiser, L., Rhee, S. Y., Sachs, M. M., Schaeffer, M., et al. (2005). Plant ontology (po): a controlled vocabulary of plant structures and growth stages. Comparative and functional genomics, 6(7-8):388–397.

Jelodar, H., Wang, Y., Yuan, C., Feng, X., Jiang, X., Li, Y., and Zhao, L. (2019). Latent dirichlet allocation (lda) and topic modeling: models, applications, a survey. Multimedia Tools and Applications, 78(11):15169–15211.

Kanehisa, M. et al. (2002). The kegg database. In Novartis Foundation Symposium, pages 91–100. Wiley Online Library.

Lau, J. H. and Baldwin, T. (2016). An empirical evaluation of doc2vec with practical insights into document embedding generation. arXiv preprint arXiv:1607.05368.

Le, Q. V. and Mikolov, T. (2014). Distributed representations of sentences and documents. corr abs/1405.4053 (2014). arXiv preprint arXiv:1405.4053.

Lee, D. D. and Seung, H. S. (1999). Learning the parts of objects by non-negative matrix factorization. Nature, 401(6755):788–791.

Lee, J., Yoon, W., Kim, S., Kim, D., Kim, S., So, C. H., and Kang, J. (2020). Biobert: a pre-trained biomedical language representation model for biomedical text mining. Bioinformatics, 36(4):1234–1240.

Li, T., Zhang, W., Yang, H., Dong, Q., Ren, J., Fan, H., Zhang, X., and Zhou, Y. (2019). Comparative transcriptome analysis reveals differentially expressed genes related to the tissue-specific accumulation of anthocyanins in pericarp and aleurone layer for maize. Scientific reports, 9(1):1–12.

Lloyd, J. and Meinke, D. (2012). A comprehensive dataset of genes with a loss-of-function mutant phenotype in arabidopsis. Plant physiology, 158(3):1115–1129.

Mikolov, T., Chen, K., Corrado, G., and Dean, J. (2013). Efficient estimation of word representations in vector space. arXiv preprint arXiv:1301.3781.

Moen, S. and Ananiadou, T. S. S. (2013). Distributional semantics resources for biomedical text processing. Proceedings of LBM, pages 39–44.

Mungall, C. J., McMurry, J. A., Köhler, S., Balhoff, J. P., Borromeo, C., Brush, M., Carbon, S., Conlin, T., Dunn, N., Engelstad, M., et al. (2017). The monarch initiative: an integrative data and analytic platform connecting phenotypes to genotypes across species. Nucleic acids research, 45(D1):D712–D722.

Oellrich, A., Walls, R. L., Cannon, E. K., Cannon, S. B., Cooper, L., Gardiner, J., Gkoutos, G. V., Harper, L., He, M., Hoehndorf, R., et al. (2015). An ontology approach to comparative phenomics in plants. Plant methods, 11(1):1–15.

OpenAI (2023). Introducing ChatGPT.

Pontes, E. L., Huet, S., Torres-Moreno, J.-M., and Linhares, A. C. (2016). Automatic text summarization with a reduced vocabulary using continuous space vectors. In International Conference on Applications of Natural Language to Information Systems, pages 440–446. Springer.

Portwood, J. L., Woodhouse, M. R., Cannon, E. K., Gardiner, J. M., Harper, L. C., Schaeffer, M. L., Walsh, J. R., Sen, T. Z., Cho, K. T., Schott, D. A., et al. (2019). Maizegdb 2018: the maize multi-genome genetics and genomics database. Nucleic acids research, 47(D1):D1146–D1154.

Radford, A., Narasimhan, K., Salimans, T., and Sutskever, I. (2018). Improving language understanding by generative pre-training.

Rehurek, R. and Sojka, P. (2010). Software framework for topic modelling with large corpora. In In Proceedings of the LREC 2010 Workshop on New Challenges for NLP Frameworks. Citeseer.

Robinson, P. N., Kö hler, S., Bauer, S., Seelow, D., Horn, D., and Mundlos, S. (2008). The human phenotype ontology: a tool for annotating and analyzing human hereditary disease. The American Journal of Human Genetics, 83(5):610–615.

Schläpfer, P., Zhang, P., Wang, C., Kim, T., Banf, M., Chae, L., Dreher, K., Chavali, A. K., Nilo-Poyanco, R., Bernard, T., et al. (2017). Genome-wide prediction of metabolic enzymes, pathways, and gene clusters in plants. Plant physiology, 173(4):2041–2059.

Schriml, L. M., Arze, C., Nadendla, S., Chang, Y.-W. W., Mazaitis, M., Felix, V., Feng, G., and Kibbe, W. A. (2012). Disease ontology: a backbone for disease semantic integration. Nucleic acids research, 40(D1):D940–D946.

Soğ ancıoğ lu, G., Öztürk, H., and Özgür, A. (2017). Biosses: a semantic sentence similarity estimation system for the biomedical domain. Bioinformatics, 33(14):i49–i58.

Szklarczyk, D., Morris, J. H., Cook, H., Kuhn, M., Wyder, S., Simonovic, M., Santos, A., Doncheva, N. T., Roth, A., Bork, P., et al. (2016). The string database in 2017: quality-controlled protein–protein association networks, made broadly accessible. Nucleic acids research, page gkw937.

Thomas, P. D., Kejariwal, A., Campbell, M. J., Mi, H., Diemer, K., Guo, N., Ladunga, I., Ulitsky-Lazareva, B., Muruganujan, A., Rabkin, S., et al. (2003). Panther: a browsable database of gene products organized by biological function, using curated protein family and subfamily classification. Nucleic acids research, 31(1):334–341.

Tseytlin, E., Mitchell, K., Legowski, E., Corrigan, J., Chavan, G., and Jacobson, R. S. (2016). Noble– flexible concept recognition for large-scale biomedical natural language processing. BMC bioinformatics, 17(1):32.

Vaswani, A., Shazeer, N., Parmar, N., Uszkoreit, J., Jones, L., Gomez, A. N., Kaiser, Ł., and Polosukhin, I. (2017). Attention is all you need. In Advances in neural information processing systems, pages 5998–6008.

Wimalanathan, K., Friedberg, I., Andorf, C. M., and Lawrence-Dill, C. J. (2018). Maize go annotation—methods, evaluation, and review (maize-gamer). Plant Direct, 2(4):e00052.

Wolf, T., Chaumond, J., Debut, L., Sanh, V., Delangue, C., Moi, A., Cistac, P., Funtowicz, M., Davison, J., Shleifer, S., et al. (2020). Transformers: State-of-the-art natural language processing. In Proceedings of the 2020 Conference on Empirical Methods in Natural Language Processing: System Demonstrations, pages 38–45.

Yanarella, C. F., Braun, I. R., Lawrence-Dill, C. J., et al. (2020). Computing on phenotypic descriptions for candidate gene discovery and crop improvement. Plant Phenomics, 2020:1963251.

Zhou, N., Jiang, Y., Bergquist, T. R., Lee, A. J., Kacsoh, B. Z., Crocker, A. W., Lewis, K. A., Georghiou, G., Nguyen, H. N., Hamid, M. N., et al. (2019). The cafa challenge reports improved protein function prediction and new functional annotations for hundreds of genes through experimental screens. Genome biology, 20(1):1–23.

