## Supplemental Table 1 for "The Case for Retaining Natural Language Descriptions of Phenotypes in Plant Databases and a Web Application as Proof of Concept"

**Table S1.** Comparing $F_1$ scores for associations and orthologous gene pair relationships.

| Approach | Category | Concat | Associations ($F_1$) | | Orthologs ($F_1$) | |
|---|---|---|---|---|---|---|
| | | | All Genes | Curated | All Genes | Curated |
| Baseline | Baseline | Yes | 0.031 | 0.041 | 0.001 | 0.001 |
| TF-IDF (Unigrams) | TF-IDF | Yes | 0.049 | 0.068 | 0.010 | 0.061 |
| TF-IDF (Unigrams & Bigrams) | TF-IDF | Yes | 0.051 | 0.072 | 0.016 | 0.054 |
| TF-IDF (Plant Article Unigrams) | TF-IDF | Yes | 0.048 | 0.067 | 0.008 | 0.057 |
| NOBLE Coder (Precise) | Annotation | Yes | 0.037 | 0.049 | 0.012 | 0.022 |
| NOBLE Coder (Partial) | Annotation | Yes | 0.042 | 0.060 | 0.003 | 0.016 |
| LDA (50 Topics) | Topic Modeling | Yes | 0.039 | 0.053 | 0.002 | 0.005 |
| LDA (100 Topics) | Topic Modeling | Yes | 0.038 | 0.053 | 0.005 | 0.004 |
| NMF (50 Topics) | Topic Modeling | Yes | 0.042 | 0.060 | 0.007 | 0.013 |
| NMF (100 Topics) | Topic Modeling | Yes | 0.043 | 0.061 | 0.006 | 0.020 |
| Doc2Vec (Wikipedia) | ML (Embeddings) | Yes | 0.033 | 0.047 | 0.015 | 0.029 |
| Doc2Vec (Plants) | ML (Embeddings) | Yes | 0.031 | 0.041 | 0.001 | 0.007 |
| Word2Vec (Wikipedia) | ML (Embeddings) | Yes | 0.042 | 0.059 | 0.003 | 0.003 |
| Word2Vec (PubMed) | ML (Embeddings) | Yes | 0.042 | 0.060 | 0.006 | 0.007 |
| Word2Vec (Plants) | ML (Embeddings) | Yes | 0.052 | 0.070 | 0.012 | 0.065 |
| BERT | ML (Embeddings) | Yes | 0.045 | 0.059 | 0.003 | 0.003 |
| BioBERT | ML (Embeddings) | Yes | 0.046 | 0.062 | 0.009 | 0.020 |
| Word2Vec (Wikipedia) | ML (Word Replacement) | Yes | 0.046 | 0.065 | 0.006 | 0.029 |
| Word2Vec (PubMed) | ML (Word Replacement) | Yes | 0.048 | 0.066 | 0.023 | 0.133 |
| Word2Vec (Plant Phenotypes) | ML (Word Replacement) | Yes | 0.048 | 0.069 | 0.018 | 0.080 |
| Baseline | Baseline | No | 0.044 | 0.067 | 0.004 | 0.008 |
| TF-IDF (Unigrams) | TF-IDF | No | 0.051 | 0.072 | 0.006 | 0.008 |
| TF-IDF (Unigrams & Bigrams) | TF-IDF | No | 0.051 | 0.072 | 0.005 | 0.007 |
| TF-IDF (Plant Article Unigrams) | TF-IDF | No | 0.051 | 0.073 | 0.004 | 0.007 |
| NOBLE Coder (Precise) | Annotation | No | 0.051 | 0.068 | 0.004 | 0.003 |
| NOBLE Coder (Partial) | Annotation | No | 0.048 | 0.069 | 0.004 | 0.005 |
| NMF (50 Topics) | Topic Modeling | No | 0.049 | 0.070 | 0.004 | 0.007 |
| NMF (100 Topics) | Topic Modeling | No | 0.049 | 0.071 | 0.005 | 0.006 |
| LDA (50 Topics) | Topic Modeling | No | 0.047 | 0.069 | 0.005 | 0.006 |
| LDA (100 Topics) | Topic Modeling | No | 0.048 | 0.069 | 0.006 | 0.008 |
| Doc2Vec (Wikipedia) | ML (Embeddings) | No | 0.051 | 0.071 | 0.007 | 0.008 |
| Doc2Vec (Plants) | ML (Embeddings) | No | 0.051 | 0.071 | 0.006 | 0.010 |
| Word2Vec (Wikipedia) | ML (Embeddings) | No | 0.049 | 0.070 | 0.006 | 0.005 |
| Word2Vec (PubMed) | ML (Embeddings) | No | 0.048 | 0.071 | 0.007 | 0.009 |
| Word2Vec (Plants) | ML (Embeddings) | No | 0.053 | 0.074 | 0.006 | 0.008 |
| BERT | ML (Embeddings) | No | 0.048 | 0.070 | 0.005 | 0.008 |
| BioBERT | ML (Embeddings) | No | 0.048 | 0.071 | 0.005 | 0.008 |
| Word2Vec (Wikipedia) | ML (Word Replacement) | No | 0.052 | 0.073 | 0.005 | 0.007 |
| Word2Vec (PubMed) | ML (Word Replacement) | No | 0.052 | 0.073 | 0.005 | 0.006 |
| Word2Vec (Plant Phenotypes) | ML (Word Replacement) | No | 0.050 | 0.072 | 0.006 | 0.007 |
| GO | Curation | | | 0.094 | | 0.059 |
| PO | Curation | | | 0.048 | | 0.001 |
| EQs | Curation | | | 0.063 | | 0.014 |
