## Supplemental Table 2 for "The Case for Retaining Natural Language Descriptions of Phenotypes in Plant Databases and a Web Application as Proof of Concept"

**Table S2.** Comparing $F_1$ scores for pathways for intraspecies and interspecies gene pairs.

| Approach | Category | Concat | Pathways, All Genes ($F_1$) | | Pathways, Curated ($F_1$) | |
|---|---|---|---|---|---|---|
| | | | Intraspecies | Interspecies | Intraspecies | Interspecies |
| Baseline | Baseline | Yes | 0.053 | 0.051 | 0.054 | 0.049 |
| TF-IDF (Unigrams) | TF-IDF | Yes | 0.107 | 0.067 | 0.116 | 0.094 |
| TF-IDF (Unigrams & Bigrams) | TF-IDF | Yes | 0.111 | 0.069 | 0.123 | 0.097 |
| TF-IDF (Plant Article Unigrams) | TF-IDF | Yes | 0.102 | 0.067 | 0.112 | 0.092 |
| NOBLE Coder (Precise) | Annotation | Yes | 0.078 | 0.055 | 0.082 | 0.073 |
| NOBLE Coder (Partial) | Annotation | Yes | 0.088 | 0.058 | 0.097 | 0.072 |
| LDA (50 Topics) | Topic Modeling | Yes | 0.084 | 0.060 | 0.089 | 0.076 |
| LDA (100 Topics) | Topic Modeling | Yes | 0.078 | 0.060 | 0.086 | 0.065 |
| NMF (50 Topics) | Topic Modeling | Yes | 0.092 | 0.073 | 0.104 | 0.097 |
| NMF (100 Topics) | Topic Modeling | Yes | 0.091 | 0.068 | 0.104 | 0.081 |
| Doc2Vec (Wikipedia) | ML (Embeddings) | Yes | 0.069 | 0.051 | 0.070 | 0.062 |
| Doc2Vec (Plants) | ML (Embeddings) | Yes | 0.060 | 0.055 | 0.062 | 0.049 |
| Word2Vec (Wikipedia) | ML (Embeddings) | Yes | 0.089 | 0.053 | 0.102 | 0.067 |
| Word2Vec (PubMed) | ML (Embeddings) | Yes | 0.095 | 0.056 | 0.114 | 0.074 |
| Word2Vec (Plants) | ML (Embeddings) | Yes | 0.105 | 0.071 | 0.115 | 0.107 |
| BERT | ML (Embeddings) | Yes | 0.087 | 0.052 | 0.102 | 0.059 |
| BioBERT | ML (Embeddings) | Yes | 0.089 | 0.051 | 0.104 | 0.060 |
| Word2Vec (Wikipedia) | ML (Word Replacement) | Yes | 0.100 | 0.063 | 0.110 | 0.088 |
| Word2Vec (PubMed) | ML (Word Replacement) | Yes | 0.105 | 0.062 | 0.114 | 0.088 |
| Word2Vec (Plant Phenotypes) | ML (Word Replacement) | Yes | 0.106 | 0.071 | 0.115 | 0.108 |
| Baseline | Baseline | No | 0.091 | 0.051 | 0.101 | 0.049 |
| TF-IDF (Unigrams) | TF-IDF | No | 0.102 | 0.067 | 0.109 | 0.099 |
| TF-IDF (Unigrams & Bigrams) | TF-IDF | No | 0.102 | 0.069 | 0.109 | 0.093 |
| TF-IDF (Plant Article Unigrams) | TF-IDF | No | 0.100 | 0.066 | 0.107 | 0.098 |
| NOBLE Coder (Precise) | Annotation | No | 0.093 | 0.057 | 0.106 | 0.081 |
| NOBLE Coder (Partial) | Annotation | No | 0.096 | 0.058 | 0.105 | 0.069 |
| NMF (50 Topics) | Topic Modeling | No | 0.094 | 0.054 | 0.102 | 0.070 |
| NMF (100 Topics) | Topic Modeling | No | 0.093 | 0.058 | 0.103 | 0.070 |
| LDA (50 Topics) | Topic Modeling | No | 0.091 | 0.056 | 0.102 | 0.069 |
| LDA (100 Topics) | Topic Modeling | No | 0.098 | 0.070 | 0.107 | 0.077 |
| Doc2Vec (Wikipedia) | ML (Embeddings) | No | 0.103 | 0.056 | 0.110 | 0.070 |
| Doc2Vec (Plants) | ML (Embeddings) | No | 0.101 | 0.063 | 0.106 | 0.077 |
| Word2Vec (Wikipedia) | ML (Embeddings) | No | 0.100 | 0.055 | 0.108 | 0.069 |
| Word2Vec (PubMed) | ML (Embeddings) | No | 0.103 | 0.060 | 0.112 | 0.082 |
| Word2Vec (Plants) | ML (Embeddings) | No | 0.106 | 0.072 | 0.113 | 0.104 |
| BERT | ML (Embeddings) | No | 0.104 | 0.057 | 0.115 | 0.069 |
| BioBERT | ML (Embeddings) | No | 0.106 | 0.057 | 0.116 | 0.079 |
| Word2Vec (Wikipedia) | ML (Word Replacement) | No | 0.104 | 0.070 | 0.112 | 0.102 |
| Word2Vec (PubMed) | ML (Word Replacement) | No | 0.104 | 0.064 | 0.111 | 0.090 |
| Word2Vec (Plant Phenotypes) | ML (Word Replacement) | No | 0.103 | 0.073 | 0.108 | 0.108 |
| GO | Curation | | | | 0.137 | 0.191 |
| PO | Curation | | | | 0.057 | 0.107 |
| EQs | Curation | | | | 0.097 | 0.049 |
